# Pink1-mediated mitophagy in the endothelium releases proteins encoded by mitochondrial DNA and activates neutrophil responses

**DOI:** 10.1101/2022.08.07.503084

**Authors:** Priyanka Gajwani, Li Wang, Shubhi Srivastava, Zijing Ye, Young-Mee Kim, Sarah Krantz, Dong-Mei Wang, Chinnaswamy Tiruppathi, Peter T. Toth, Jalees Rehman

**Affiliations:** Department of Pharmacology and Regenerative Medicine, University of Illinois College of Medicine, Chicago, IL 60612, USA; Division of Cardiology, Department of Medicine, University of Illinois College of Medicine, Chicago, IL 60612, USA; Department of Biochemistry and Molecular Genetics, The University of Illinois College of Medicine, Chicago, IL 60607; Research Resources Core, University of Illinois at Chicago, Chicago, IL 60612, USA; University of Illinois Cancer Center, Chicago, IL 60612, USA

**Author notes:** Please address correspondence to: Jalees Rehman, Department of Biochemistry and Molecular Genetics, The University of Illinois College of Medicine 900 South Ashland Ave (MC 669), Chicago, IL, 60607. **Author Contributions:** The study was conceived and supervised by J.R. The experiments were designed, performed, and analyzed by P.G., L.W., S.S., Z.Y, Y.M.K., D.M.W, S.K., C.T., P.T.T. and J.R. The initial manuscript draft was written by P.G. and J.R. All authors reviewed the manuscript and provided feedback and revisions.

**Keywords:** Mitophagy, Mitochondria, Inflammation, Endothelial cells, Formylated proteins, Formyl Peptide Receptors, Neutrophils, Host defense

## Abstract

Given their ancient evolutionary origins, eukaryotic mitochondria possess multiple vestiges of their prokaryotic ancestors. One such factor is the N-terminal formylation of proteins encoded by mitochondrial DNA. N-formylated proteins are also released by bacteria and trigger activation of immune cells such as neutrophils. Growing evidence indicate that circulating levels of mitochondrial formyl proteins are elevated in the serum of patients with excessive inflammatory responses and trigger neutrophil activation like their bacterial counterparts. However, the cellular source of these proteins, and the mechanism by which they are released into the circulation is not known. In this study, we have identified vascular endothelial cells as a source of mitophagy induced release of formyl proteins in response to inflammatory mediators in vitro. Mechanistically, endothelial mitophagy required activation of the Pink1 pathway. Using liposomal delivery of sgRNA targeting Pink1 in mice expressing endothelial-specific Cas9, we developed a mouse model in which Pink1 is specifically depleted in the endothelium. Deletion of endothelial Pink1 was remarkably protective in endotoxin-induced lung inflammation, resulting in reduced neutrophil infiltration and significantly reduced death in mice. We thus propose that endothelial cells upregulate pro-inflammatory mitophagy in response to inflammation, leading to release of mitochondrial formyl peptides and detrimental neutrophil recruitment into the lung.

## Introduction

Vascular endothelial cells lining the blood vessels are the first point of contact for circulating immune cells that transmigrate into a tissue and therefore endothelial cells play a critical role in regulating immune response(Kolaczkowska and Kubes 2013, Amersfoort, Eelen et al. 2022). Through the modulation of vascular permeability, expression of surface markers and secretion of signaling factors, endothelial cells recruit and direct immune cells such as neutrophils to infected tissues (Pober and Sessa 2007, Muller 2016, Al-Soudi, Kaaij et al. 2017, Filippi 2019). Thus, endothelial function is an important determinant of the rate and extent of inflammatory activation. Recent studies of endothelial function suggest a critical role for endothelial metabolism driving endothelial migration and angiogenesis (De Bock, Georgiadou et al. 2013, De Bock, Georgiadou et al. 2013). The role of mitochondria in endothelial function is less clear, primarily because endothelial metabolism studies have focused on glycolytic pathways which predominantly drive ATP production in endothelial cells (Davidson and Duchen 2007, De Bock, Georgiadou et al. 2013, De Bock, Georgiadou et al. 2013). However, beyond ATP production, endothelial mitochondria serve as important signaling organelles, through the control of ROS, NO and Ca^2+^ signaling(Quintero, Colombo et al. 2006, Kluge, Fetterman et al. 2013, Tiku, Tan et al. 2020). Endothelial mitochondria undergo depolarization in response to inflammatory mediators such as the cytokine TNFα (Chen, Reece et al. 1999, Corda, Laplace et al. 2001) but the underlying molecular mechanisms and impact on host defense need to be defined.

In response to depolarization, damaged mitochondria are typically sequestered away from the rest of the mitochondrial pool and targeted to the lysosomes for degradation through the process of mitochondrial autophagy referred to as mitophagy(Onishi, Yamano et al. 2021). Mitophagy most often occurs through the Pink1/Parkin pathway, which results in the ubiquitination of damaged mitochondria that are then engulfed by an autophagosome and transported to the lysosome(Palikaras, Lionaki et al. 2018, Ng, Wai et al. 2021). Intriguingly, the global deletion of Parkin leads to reduced endothelial inflammatory activation suggesting a pro-inflammatory role for mitophagy (Letsiou, Sammani et al. 2017). Several recent studies have suggested that mitochondria and mitochondrial damage associated molecular patterns (DAMPs) are actively released by cells in response to inflammatory and other stimuli(Zhang, Raoof et al. 2010, Dorward, Lucas et al. 2017, D’Acunzo, Pérez-González et al. 2021), and can promote pro-inflammatory responses (Puhm, Afonyushkin et al. 2019).

Given their endosymbiont evolutionary origin, mitochondria contain several remnants of their prokaryotic ancestors which may trigger immune responses in mammalian organisms (Zhang, Raoof et al. 2010). Proteins encoded by mitochondrial DNA are translated in the mitochondria by ribosomes that resemble prokaryotic translation machinery and differ from their nuclear-encoded counterparts because the initiating methionine contains an additional N-formyl group (Tucker, Hershman et al. 2011). Bacterial peptides and proteins contain N-formyl groups which are recognized by mammalian immune cells and initiate inflammatory activation (Bloes, Kretschmer et al. 2015), but interestingly endogenous mitochondrial formylated proteins that are released by mammalian cells can also bind to formyl peptide receptors on the surface of innate immune cells, inducing activation and transmigration(Rongvaux 2018). While this phenomenon has been long noted in sterile injury due to physical trauma(McDonald, Pittman et al. 2010), mitochondrial formylated proteins have also been observed in the serum of patients with sepsis which are characterized by excessive immune responses, suggesting that mitochondrial formyl-protein release may also occur during inflammatory injury(Wenceslau, McCarthy et al. 2015, Kwon, Suh et al. 2021, Yuan, Zeng et al. 2021), although it is not known which cell types release these DAMPs and which signaling pathways trigger the release.

In this study, we observed that endothelial cells in the lung upregulate mitophagy in response to the systemic delivery of the bacterial endotoxin lipopolysaccharide (LPS) *in vivo*. The inflammatory mediator TNFα, which is released by immune cells in response to LPS induces mitophagy through the Pink1/Parkin pathway. Mice with endothelial-specific depletion of Pink1 using targeted CRISPR/Cas9 editing in mouse lungs using liposomal delivery exhibited improved survival to LPS-induced endotoxemia and reduced neutrophil invasion into the lungs. These data suggest that endothelial Pink1-mediated mitophagy acts as a pro-inflammatory amplification pathway that could be targeted to reduce excessive inflammatory responses.

## Results

### Endotoxemic inflammation induces mitophagy in lung vascular endothelial cells

Since inflammation induces mitochondrial depolarization in endothelial cells(Corda, Laplace et al. 2001), we first sought to determine whether inflammation induced mitophagy in the vasculature in vivo. We evaluated in vivo mitophagy using the mitophagy reporter Mitokeima mice(Sun, Malide et al. 2017). Mitokeima mice globally express the biosensor Mitokeima, which distinguishes between cytosolic mitochondria at neutral pH and lysosomal mitochondria, “mitolysosomes”, that are at an acidic pH. To label the vascular system, mice were retro-orbitally injected with a fluorescently labeled lectin that specifically binds to mouse endothelial cells - Isolectin-B4 conjugated to DyLight 647. Ex vivo imaging by confocal microscopy revealed regions of high and low mitophagy in the whole lung tissue **(Figure 1A, Figure 1 – figure supplement 1**). To distinguish between endothelial and non-endothelial mitophagy, we generated an image analysis “mask” based on the endothelial Isolectin-B4 channel and applied it to determine the ratio of Acidic:Neutral mitokeima. A 3D reconstruction of lung vascular mitophagy generated using Imaris image analysis software showed that this method accurately isolates the ratiometric Mitokeima signal from the endothelium, while excluding the surrounding tissue (**Figure 1B**).

**Figure 1:**
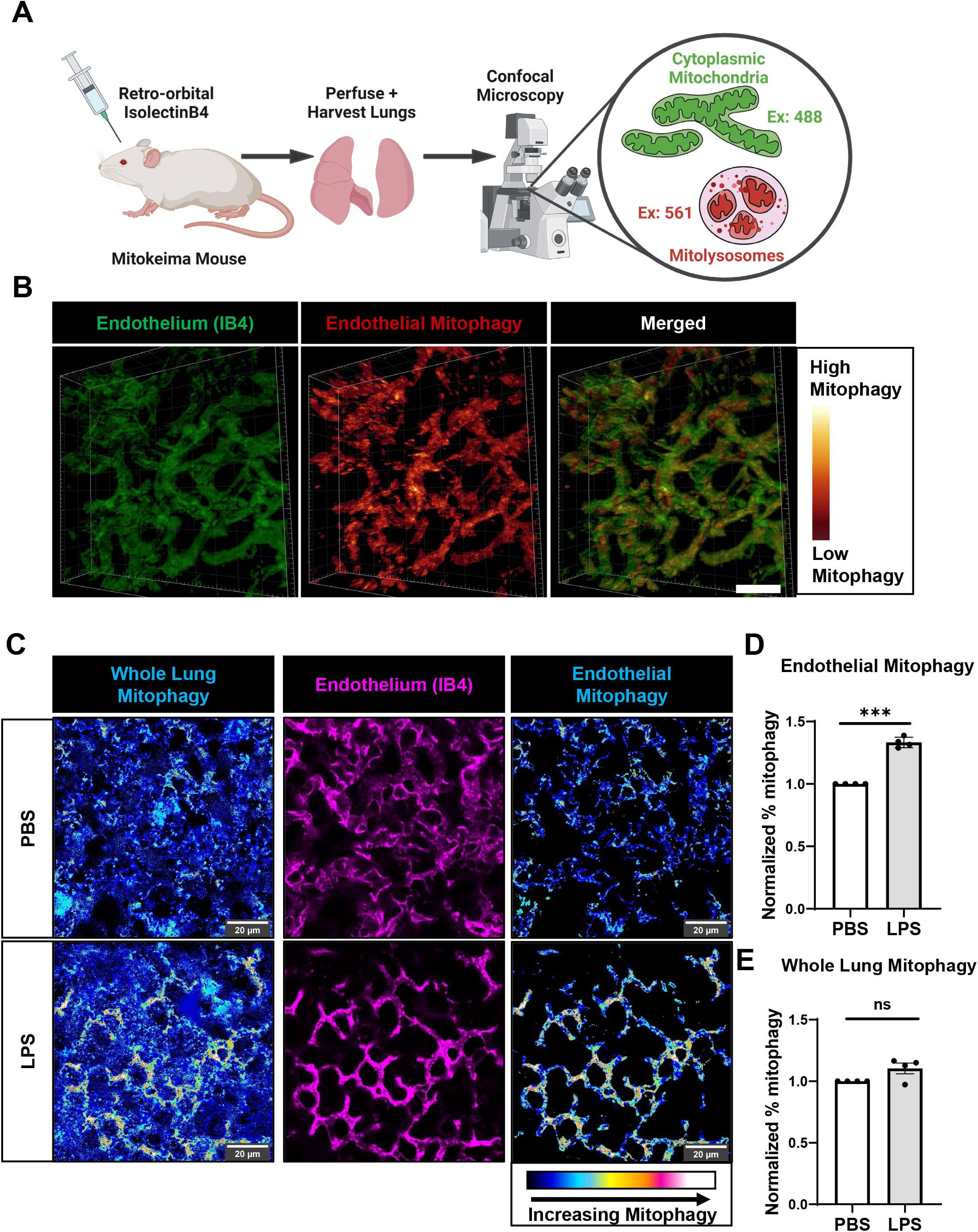
Lung vascular endothelial cells initiate mitophagy in response to endotoxemic inflammation. Mice expressing the mitophagy biosensor Mitokeima were injected with Isolectin B4, to label endothelial cells. Lungs were harvested and perfused, and mitophagy was visualized in the whole, unsectioned lung by confocal microscopy **(A)**. A 3D mask of the endothelium was constructed and used to isolate the Mitokeima acidic/neutral ratio specifically in endothelial cells **(B)**. Scale Bar: 10μm. Using this method, mitophagy was measured in an endotoxemia model of inflammation. Lungs from Mitokeima mice were visualized 6 hours post i.p. LPS injection (8mg/kg) **(C)**. Endothelial **(D)** and whole lung **(E)** mitophagy was measured by calculating the ratio of acidic to neutral Mitokeima (n=4 mice, 10-20 fields of view per mouse). Statistical significance between PBS and LPS treated mice was evaluated by t-test Source data for (D) and (E) available in Figure 1 Source Data 1.

We applied this method to quantify mitophagy in the lungs of mice injected with the bacterial endotoxin Lipopolysaccharide (LPS). Mitokeima mice were injected with LPS (8mg/kg, i.p.) to induce endotoxemic inflammation, and lungs were isolated and imaged 6 hours post-LPS injection. Compared to PBS-injected control mice, LPS-injected mice had an approximately 30% increase in endothelial mitophagy (**Figure 1C, D**). Mitophagy in the whole lung did not significantly change (**Figure 1E**). Thus, these results indicate that systemic inflammation induced by LPS specifically induces mitophagy in the lung vasculature.

### Live cell imaging of mitophagy induced by the inflammatory mediator TNFα

The inflammatory mediator TNFα is secreted by monocytes in response to infection and LPS-induced inflammation and is a major regulator of inflammatory activation of endothelial cells(Sethi and Hotamisligil 2021). We thus hypothesized that TNFα was the mediator of LPS-induced endothelial mitophagy observed in vivo. Primary human lung microvascular endothelial cells (HLMVECs) were transduced to express Mitokeima and treated with TNFα over a time course of 6 hours to monitor mitophagy.

Mitophagy was quantified as the ratio of acidic mitochondria/total mitochondria as analyzed by confocal microscopy. TNFα significantly induced mitophagy in HLMVECs within 3 hours (**Figure 2A,B**).

**Figure 2:**
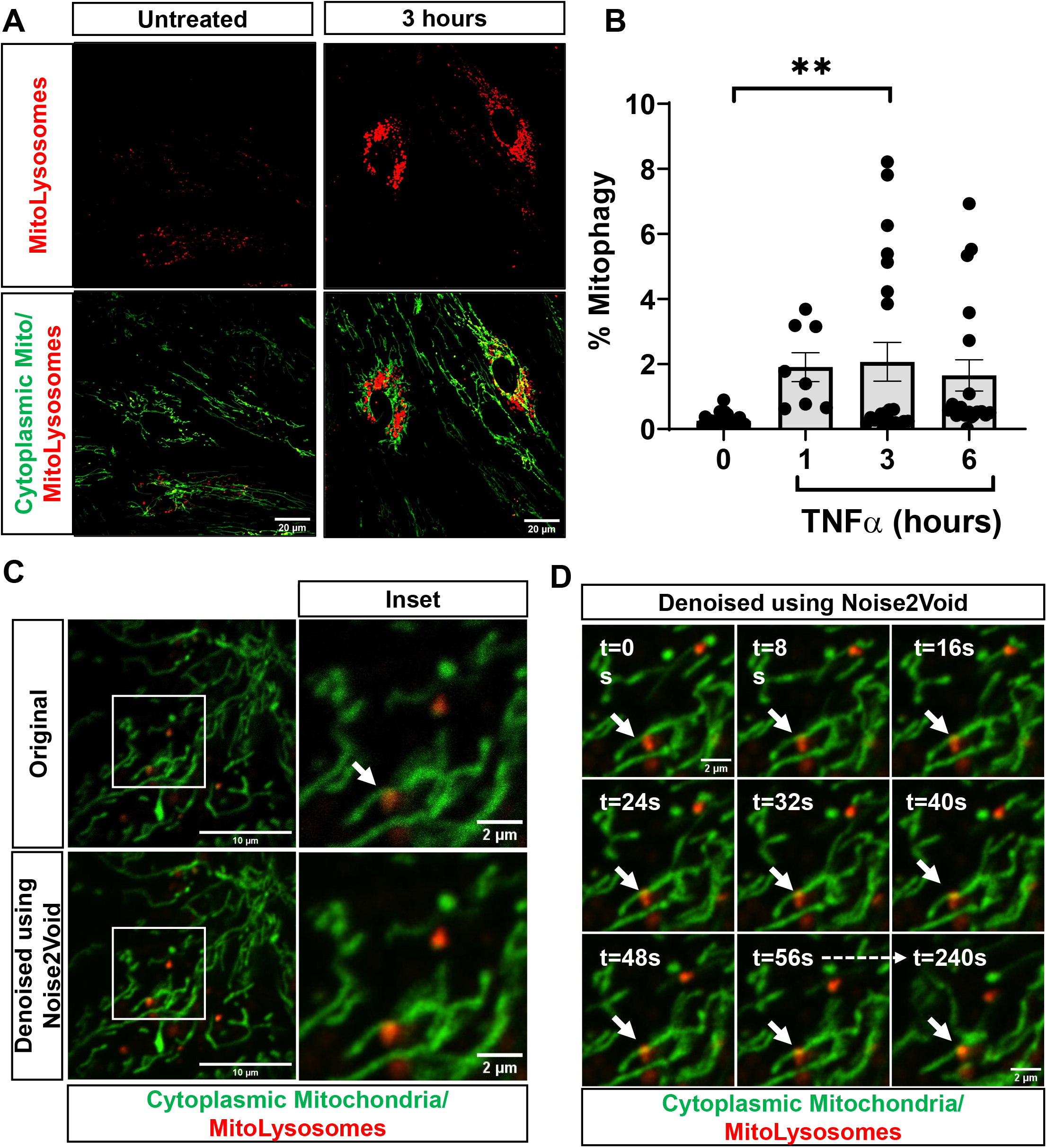
TNFα induces endothelial mitophagy within 3 hours. HLMVECs expressing mitokeima were treated with the inflammatory mediator TNFα (10ng/mL), and mitophagy visualized at 1-, 3- and 6-hours post TNFα exposure, with representative images at 0 and 3 hours **(A)**. Mitophagy was calculated by measuring the ratio of acidic mitochondria to total mitochondrial area in each visual field **(B)**. n=3 independent experiments. Source data for (B) provided in Figure 2 Source Data 1. To better visualize mitokeima in endothelial cells, images and movies were denoised using the Noise2Void denoising algorithm. Representative images of a cell treated with TNFα for 3 hours show that denoising reveals interactions between cytoplasmic mitochondria and mitolysosomes **(C)**. Interactions between cytoplasmic and lysosomal mitochondria persist over several minutes **(D)**.

To understand the dynamics of TNFα induced mitolysosomes, we imaged the time course of live TNFα treated HLMVECs expressing Mitokeima. However, due to the high degree of noise present in high resolution images of mitolysosomes, we performed image de-noising using the deep-learning algorithm Noise2Void(Krull, Buchholz et al. 2019). Noise2Void is a denoising technique which does not require an additional data set for training and can be trained on experimental images. Denoising significantly improved Mitokeima imaging, allowing the time lapse visualization of interactions between mitolysosomes and cytoplasmic mitochondria (**Figure 2C**) in HLMVECs treated with TNFα. Movies of these cells show that TNFα induced mitolysosomes make several prolonged contacts with cytoplasmic mitochondria, lasting several minutes (**Figure 2D, Supplemental Movie 1**).

### Pink1 mediates TNFα-induced endothelial mitophagy

We next sought to determine whether the Pink1/Parkin pathway, which is a major mitophagy initiating pathway in multiple cell types(McWilliams and Muqit 2017) meditated mitophagy in endothelial cells. In healthy, polarized mitochondria, Pink1 is inserted into the outer mitochondrial membrane, where it is cleaved by mitochondrial proteases, and degraded (Matsuda, Sato et al. 2010, Yamano and Youle 2013). However, when mitochondria are depolarized, Pink1 is stabilized and recruits the E3 ubiquitin ligase Parkin to ubiquitinate the outer mitochondrial membrane. Parkin mediated ubiquitination serves as the initiating signal for mitophagy(Lazarou, Sliter et al. 2015). Thus, stabilization of Pink1 indicates the activation of this pathway. HLMVECs treated with TNFα had significantly increased Pink1 protein levels within 1 hour, which was sustained over 24 hours after treatment (**Figure 3A, B**). This stabilization of Pink1 indicates that TNFα activates mitophagy through the Pink1/Parkin pathway.

**Figure 3:**
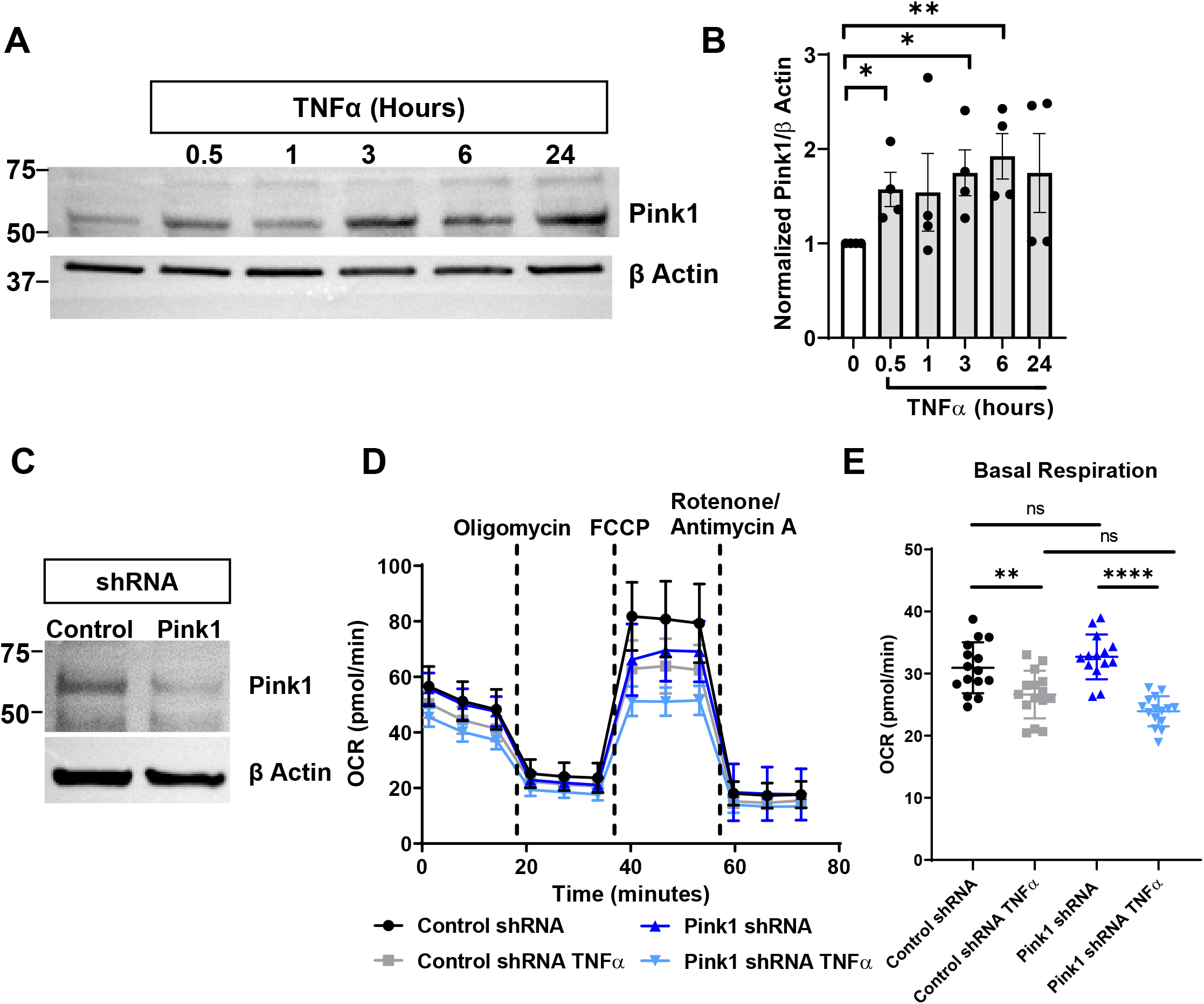
Pink1 mediates TNFα-induced endothelial mitophagy, but not mitochondrial damage. HLMVECs were treated with TNFα for 0.5-24 hours, and Pink1 protein levels were measured by western blot **(A)** and quantified **(B)**. Statistical analysis between baseline and each timepoint was done by t-test. The impact of TNFα-Pink1 mitophagy on endothelial metabolism was determined by performing a Seahorse mitochondrial stress test **(C)**, measuring the oxygen consumption rate with and without shRNA mediated deletion of Pink1 **(D)**. Basal respiration rate was plotted. Representative graphs are shown. Original western blot images provided in Figure 3 Source Data 1. Source Data for (B) (D) and (E) available in Figure 3 Source Data 2.

As mitophagy serves an important role in mitochondrial quality control, we examined whether TNFα-induced Pink1 mitophagy impacted the metabolic efficiency of endothelial mitochondria. To investigate the role of Pink-1 mediated mitophagy in mitochondrial metabolism we downregulated the cellular level of Pink-1 through shRNA (**Figure 3C**). We performed a Seahorse Analyzer mitochondrial stress test, measuring oxygen consumption as an indicator of mitochondrial oxidative phosphorylation (**Figure 3D**). In the mitochondrial stress test, HLMVECs treated with TNFα for 3 hours had reduced basal oxygen consumption, however this decrease was not dependent on Pink1 depletion (**Figure 3E**). These data suggest that TNFα-induced Pink1-mediated mitophagy does not significantly impact endothelial mitochondrial metabolism.

### Endothelial Pink1 exacerbates endotoxemia induced death

To understand the importance of endothelial Pink1-induced mitophagy in inflammation, we used a CRISPR/Cas9 approach to specifically delete endothelial Pink1 in vivo. We generated mice that express Cas9 specifically in endothelial cells by crossing knock-in Cas9 mice(Platt, Chen et al. 2014) with mice expressing Cre under the endothelial specific CDH5 (VE-Cadherin) promoter (provided by Dr. Ralf Adams). Plasmid containing sgRNA against Pink1 was encapsulated in cationic liposome, a formulation that has previously been employed to deliver genes to the lung endothelium via an intravenous route as the lung endothelium is the first microvascular bed encountered by intravenously injected liposomes (Liu, Zhang et al. 2019). Intravenous injection of Pink1 sgRNA containing liposomes led to deletion of Pink1 specifically in the Cas9 expressing endothelium (Pink1^EC-/-^ mice) (**Figure 4A**). Administration of liposome encapsulated Pink1 sgRNA resulted in an approximately 80% reduction in lung endothelial Pink1 in Cas9-expressing mice compared to age-matched wildtype C57 mice (WT mice) (**Figure 4 B,C**), but did not affect Pink1 expression in non-endothelial cells (**Figure 4D**).

**Figure 4:**
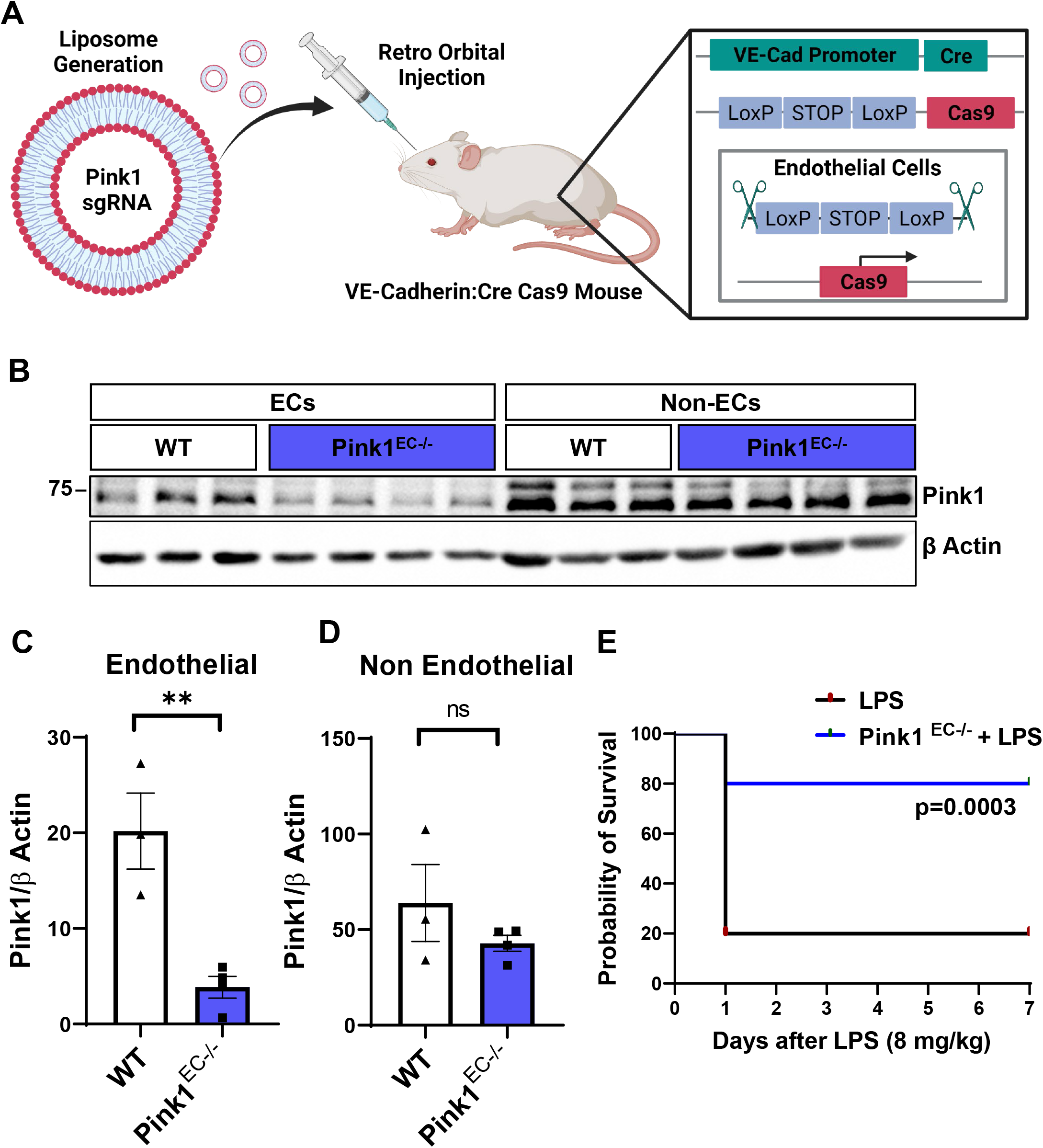
Deletion of endothelial Pink1 protects against endotoxin induced death. Mice were bred to express Cas9 in cells expressing Cre recombinase under a VE-Cadherin promoter, ensuring Cas9 expression specifically in endothelial cells. sgRNA against Pink1 was delivered to Cas9 expressing, or control C57 mice aged 8-12 weeks by retro-orbital i.v. injection of sgRNA containing liposomes **(A)**, leading to a ∼80% knockdown in endothelial cells but not in non-endothelial cells **(B-D)**. Statistical significance between Control and Pink1^EC-/-^ groups was analyzed by t-test. N=3-4 mice per group. Control and Pink1^EC-/-^ mice were injected with LPS (8mg/kg), and survival was monitored over 7 days **(E)**. n=10 male and 10 female mice per group. Uncropped western blot images for (B) available in Figure 4 Source Data 1. Source Data for (C) and (D) provided in Figure 4 Source Data 2.

To determine whether endothelial Pink1 plays a role in determining inflammatory outcome, we first examined the survival of Pink1^EC-/-^ mice in response to LPS-endotoxemia. WT and Pink1^EC-/-^ mice were injected with LPS (8mg/kg, i.p.) and survival was monitored over 7 days. Pink1^EC-/-^ mice displayed significantly improved survival compared to WT mice (**Figure 4E**). This strong pro-survival effect of EC-specific Pink1 deletion shows that endothelial Pink1 is a key mediator of pro-inflammatory activation and mortality role in LPS mediated endotoxemia. Furthermore, as shown in **Figure 4E**, the protective effect of Pink1^EC-/-^ manifested early on, with the majority of the WT mice dying on the first day following LPS administration. The early timing of this protective effect suggests that endothelial Pink1 is involved in aggravating inflammatory injury, as opposed to inhibiting pathways involved in lung regeneration.

We reasoned that endothelial Pink1 may increase inflammatory injury by increasing LPS-induced vascular permeability. To assess whether Pink1^EC-/-^ mice had altered endothelial characteristics, we determined the extent of lung edema, as measured by lung wet-to-dry weight ratio, and vascular permeability as measured by permeability to Evans Blue-conjugated Albumin (EBA), in LPS injected WT and Pink1^EC-/-^ mice. Deletion of endothelial Pink1 did not affect vascular permeability to EBA (**Figure 4 – figure supplement 1A**), nor did it affect loss of the endothelial adherens junction protein VE-Cadherin, which is an important regulator of endothelial barrier function (**Figure 4 – figure supplement 1B**). Additionally, Pink1^EC-/-^ did not alter LPS-induced lung edema (**Figure 4 – figure supplement 2**) Thus, although endothelial Pink1 mediates inflammatory lung injury, this is likely not due to direct effects on the lung vascular barrier integrity.

### Endothelial Pink1 increases neutrophil recruitment and activation in the lung

Given the strong protective effect of endothelial Pink1 deletion in inflammatory injury, and the limited effect on endothelial barrier function, we hypothesized that endothelial Pink1 induces inflammatory injury by acting on the recruitment of immune cells such as neutrophils which are key mediators of lung injury and death in endotoxemia-induced inflammatory lung injury (Hayashi, Means et al. 2003, Nathan 2006, Bachmaier, Stuart et al. 2022). Neutrophils are recruited into the lung 2-24 hours following systemic delivery of LPS where they are early drivers of inflammation induced tissue injury (Matute-Bello, Frevert et al. 2008, Zemans and Matthay 2017). We thus measured the effect of endothelial Pink1 deletion on LPS induced infiltration of neutrophils into the lung. WT and Pink1^EC-/-^ mice were injected with LPS (8mg/kg, i.p.) and lungs harvested 6 and 24 hours post-LPS injection. CD45 and Ly6G were used as markers to differentiate the neutrophil population in whole lung samples. The number of CD45+Ly6G+ neutrophils was measured by flow cytometry and normalized to the total number of cells in the lung. Pink1^EC-/-^ mice had significantly reduced neutrophil infiltration at 6 hours compared to their age-matched control counterparts **(Figure 5A,B**). Interestingly, this difference appeared only at 6 hours but did not persist at 24 hours. To further establish the importance of endothelial Pink1 in neutrophil-mediated immune response, we measured the activation of neutrophils infiltrated in lungs due to LPS induced inflammation, using CD11b as a marker for activated neutrophils. CD11b expression in CD45+Ly6G+ cells showed that neutrophil activation was compromised in Pink1^EC-/-^ mice at 24 hours (**Figure 5C,D**). These results suggest that early neutrophil recruitment is important for effective neutrophil activation, and that both processes are sensitive to Pink1^EC-/-^. The decreased early neutrophil recruitment also resulted in significantly reduced levels of the pro-inflammatory cytokine IL-1β in the lung (**Figure 5E,F**).

**Figure 5:**
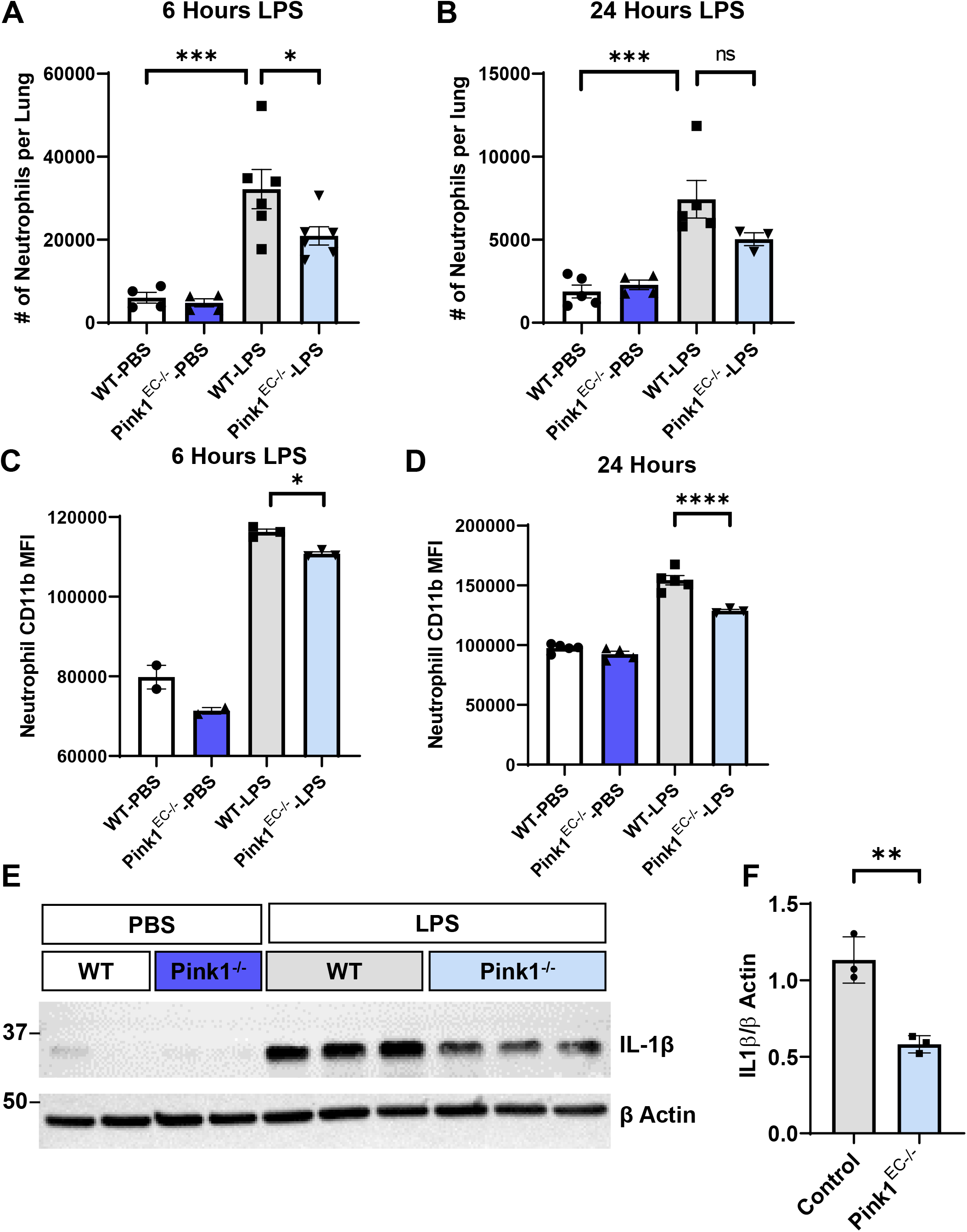
Endothelial Pink1 increases neutrophil recruitment and activation in the lung. Control (WT) and endothelial specific Pink1 knock out (Pink1^EC-/-^) mice were injected with LPS (8mg/kg). 6- and 24- hours later, lungs were perfused and harvested, and analyzed for the number of infiltrated Ly6G+ neutrophils by flow cytometry **(A,B)**. Neutrophil activation was measured by CD11b expression on Ly6G+ cells **(C,D)**. n=3-6 mice per group. IL1-β levels in the whole lung were measured by western blot **(E-F)**. n=2 mice for PBS, and n=3 mice for LPS treated groups. Statistical significance between IL-1β levels in Control and Pink1EC^-/-^ was determined by t-test. Source Data for (A), (B), (C), (D) and (F) provided in Figure 5 Source Data 1. Original western blot image for (E) available in Figure 5 Source Data 2.

One possible mechanism through which endothelial cells may alter neutrophil recruitment is through expression of the adhesion molecule ICAM-1, which is involved in neutrophil adhesion and transmigration into the lung(Yang, Froio et al. 2005). Thus, we measured ICAM-1 levels in the CD31+ endothelial cells of WT and Pink1^EC-/-^ mice injected with LPS for 6 hours. LPS induced similar activation of ICAM-1 in control and Pink1^EC-/-^ mice (**Figure 5 – figure supplement 1**), suggesting that changes in neutrophil recruitment are independent of endothelial ICAM-1 expression.

### Endothelial cells release mitochondrial formylated peptides in response to inflammation

We next examined alternative pathways through which endothelial mitochondria may interact with neutrophils. Besides interaction with adhesion molecules, Neutrophil recruitment is also heavily regulated by activation of formyl peptide receptors (FPR), which recognize bacterial proteins that contain an additional formyl group on the initiating methionine(Dorward, Lucas et al. 2015). However, given the endosymbiont origins of mitochondria and expression of N-formylated proteins by mitochondria which can activate pro-inflammatory FPRs on the surface of neutrophils, we investigated the activation of the Erk pathway which is a key signaling pathway downstream of FPR activation in neutrophils (Dorward, Lucas et al. 2015, Zhang, Liu et al. 2016, Dorward, Lucas et al. 2017). Thus, we hypothesized that endothelial mitochondria were a source of inflammation induced formyl peptide release, leading to increased neutrophil recruitment. To determine whether formyl peptides were among the factors released by endothelial cells, we looked for presence of formylated proteins in the cell culture medium. The mitochondrial protein ND6 is one of the thirteen proteins encoded in the mitochondrial DNA and is thus often used as an indicator for the presence of mitochondrial formylated proteins(Gabl, Sundqvist et al. 2018, Kwon, Suh et al. 2021). We thus performed an ELISA to measure relative ND6 levels in the cell culture medium of HLMVECs treated with TNFα or the mitochondrial uncoupler and mitophagy inducer FCCP. Both TNFα and FCCP induced a ∼30% increase in ND6 levels released by endothelial cells (**Figure 6A,B**).

**Figure 6:**
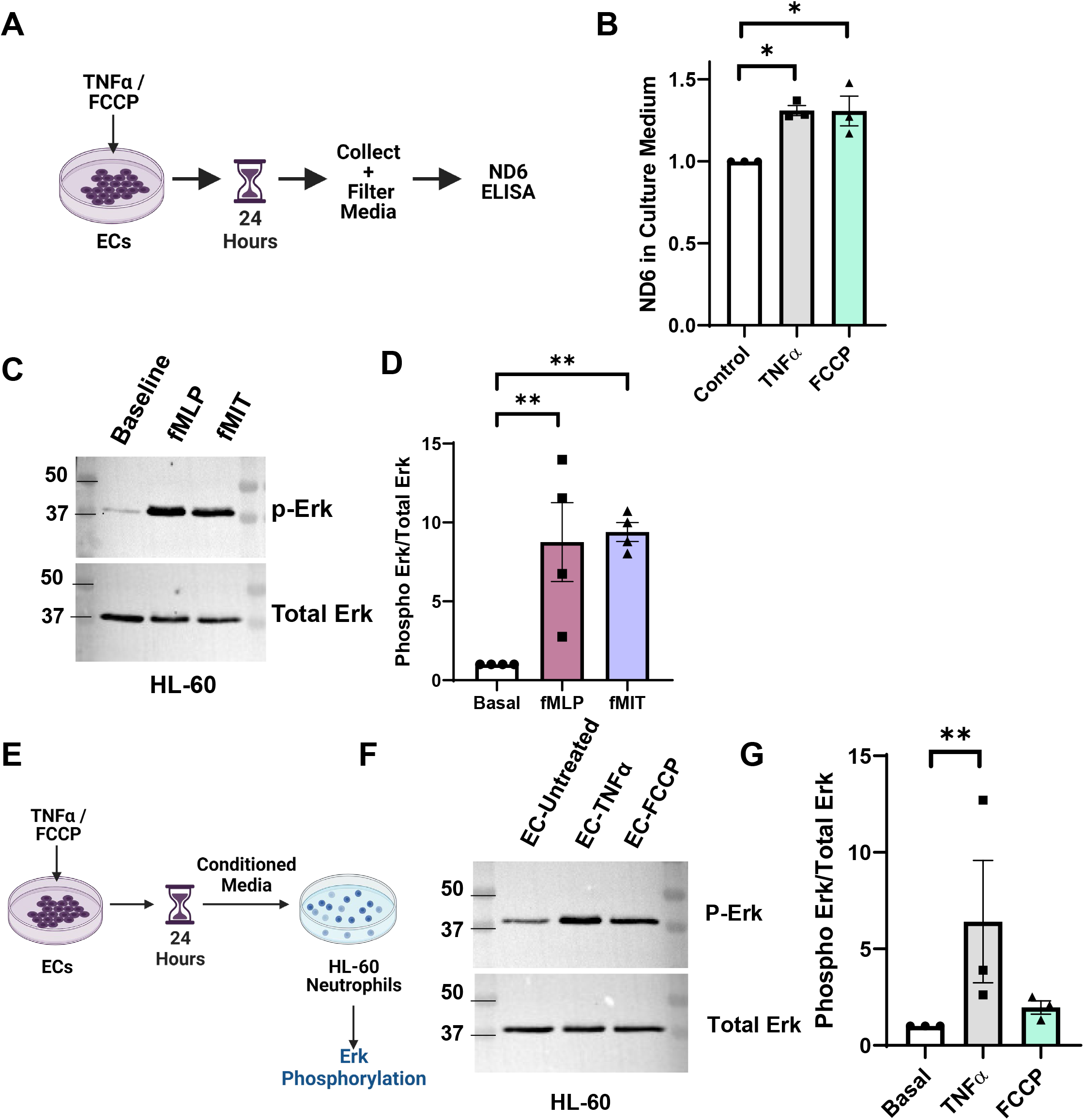
Endothelial cells release mitochondrial formylated peptides in response to inflammation and mitochondrial damage. HLMVECs were treated with TNFα or FCCP for 24 hours. Cell culture media was collected, filtered and analyzed for the presence of ND6 by ELISA **(A,B)**. n=3 independent experiments. HL60s were treated with bacterial formyl-peptide fMLP (10nM) or human mitochondrial formyl-peptide fMIT (10nM) for 10 minutes. Cells were lysed and analyzed for Erk phosphorylation by western blot **(C,D)**. n=4 independent experiments. Statistical analysis was performed by One-way ANOVA, with Holms-Sidak test for multiple comparisons. HLMVECs were treated TNFα or FCCP or DMSO for 24 hours. Media (EC-untreated, EC-TNFα, EC-FCCP) was collected and filtered to remove cell debris. Conditioned media was used to resuspend serum-starved HL-60 derived neutrophils (HL-60) for 10 minutes before cells were lysed and analyzed for phosphorylation of Erk **(E,F)**. Statistical analysis was performed using One-way ANOVA with Kruskal-Wallace test for multiple comparisons. n=3 independent experiments. Original Data for (B), (D) and (G) available in Figure 6 Source Data 1. Uncropped western blot images for (C) and (F) provided in Figure 6 Source Data 2.

To compare the ability of mitochondrial formylated peptides to activate Erk signaling in neutrophils to bacterial formylated peptides, we exposed neutrophils to purified bacterial formyl peptides (fMLP) and human mitochondrial formyl peptides (fMIT). fMIT refers to an N-formylated peptide made of the first 6 amino acids of human ND6. HL-60 derived neutrophils (referred to as HL-60) were treated with fMLP and fMIT for 10 minutes and generated an approximately equal increase in Erk phosphorylation (**Figure 6C,D**). We next determined whether factors released by activated endothelial cells led to a similar phosphorylation of Erk in neutrophils. Endothelial cells were treated with either TNFα or the mitophagy inducer FCCP. 24 hours later, cell culture medium was collected, filtered, and added to HL-60 cells for 10 minutes (**Figure 6E**). TNFα, and notably also FCCP mediated mitophagy in endothelial cells caused the release of factors into the cell culture medium that activated Erk phosphorylation in neutrophils (**Figure 6F,G**).

## Discussion

In this study, we uncovered a novel pro-inflammatory role of Pink1-mediated mitophagy in endothelial cells. Inflammatory activation induced mitophagy in endothelial cells, both in vitro and in an in vivo endotoxin model of lung injury that is characterized by excessive endothelial inflammation and influx of neutrophils (Bachmaier, Toya et al. 2007, Kolaczkowska and Kubes 2013, Zhang, Gao et al. 2022). The observed endothelial mitophagy was mediated by Pink1 activation. Deletion of Pink1 in endothelial cells in vivo resulted in increased survival in endotoxin-injected mice and significantly reduced neutrophil recruitment and activation in the lung. Lastly, we found that in response to both inflammation- and FCCP-induced mitochondrial stress, endothelial cells released ND6, a formylated mitochondrial protein and potent recruiter of neutrophils (**Figure 7**).

**Figure 7:**
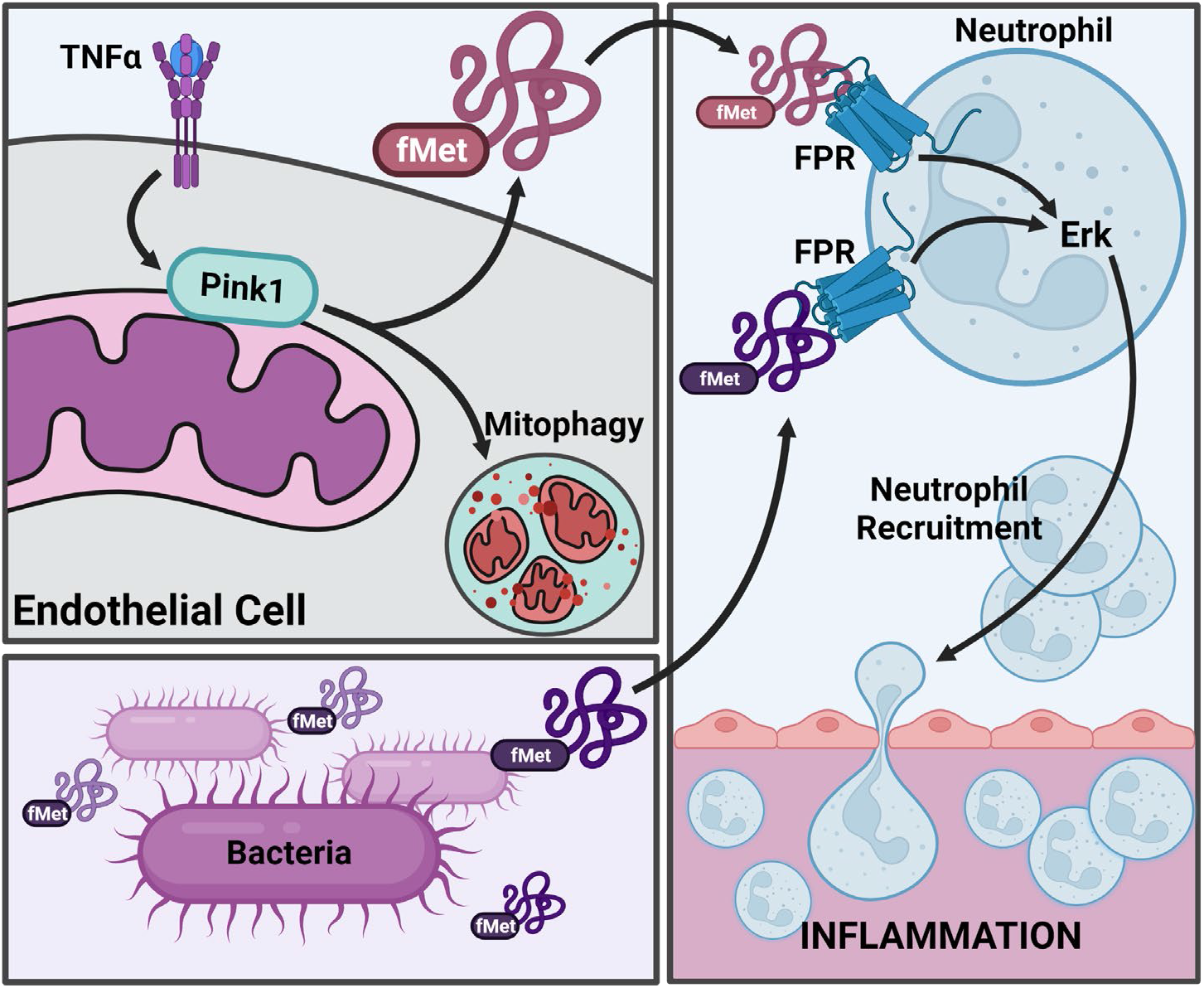
Inflammation induced endothelial mitophagy releases formylated proteins to enhance inflammation. In response to inflammatory stimulus, Pink1 is activated in endothelial cells, leading to mitophagy and release of mitochondrial proteins such as ND6 which contain a formylated methionine (fMet) at the N-terminus. Bacteria, which share a prokaryotic ancestor with mitochondria, also produce and release N-formyl proteins. Both mitochondrial and bacterial N-formyl proteins activate neutrophils through formyl peptide receptors (FPRs), leading to increased Erk phosphorylation and increased recruitment. Excessive neutrophil recruitment leads to increased aberrant inflammation.

Lung endothelial cells displayed increased mitophagy in response to inflammatory mediators both in vivo and in vitro, as visualized using the mitophagy biosensor Mitokeima. Mitokeima has been previously used to examine whole lung mitophagy, making it difficult to ascertain the role of endothelial mitophagy. Thus, to measure mitophagy specifically in the endothelium, we developed a method to isolate the endothelial Mitokeima signal from the rest of the lung in whole-organ imaging. By employing this method, we found that LPS significantly induced mitophagy in the mouse lung vascular endothelium. The inflammatory mediator TNFα, which is generated by immune cells in response to LPS, induced mitophagy in primary human lung endothelial cells. We used the deep-learning denoising algorithm Noise2Void(Krull, Buchholz et al. 2019) to allow for the visualization of interactions between mitochondria in the endothelial cytoplasm with mitochondria-containing lysosomes, suggesting that TNFα induces stable mitochondrial-lysosome contacts. These inter-organellar contacts have been implicated in transferring metabolites, altering Ca^2+^ signaling, and inducing mitochondrial fission(Wong, Ysselstein et al. 2018, Wong, Kim et al. 2019, Peng, Wong et al. 2020).

Using an endothelial-specific Cas9 mouse, and lysosomal delivery of Pink1 sgRNA, we were able to generate mice that lacked Pink1 specifically in endothelial cells, circumventing the confounding effects of global mitophagy knockouts. A surprising finding of this study was that deletion of Pink1 in endothelial cells drastically reduced endotoxin-induced death in mice, indicating a pro-inflammatory role of endothelial Pink1. By contrast, mitophagy in other immune cells is typically associated with reduced inflammatory activation(Harris, Deen et al. 2018, Sliter, Martinez et al. 2018), such as in macrophages where mitophagy is linked to reduced inflammasome activation and IL-1β production(Zhong, Umemura et al. 2016). Thus, mitophagy appears to have uniquely detrimental consequences in inflamed endothelial cells. This pro-inflammatory cost of endothelial mitophagy may thus provide an explanation as to why endothelial cells have a relatively small share of mitochondria compared to other cell types, with mitochondria taking up only 2-3% of endothelial cytoplasm(Kluge, Fetterman et al. 2013). Deletion of endothelial Pink1 significantly reduced LPS-induced neutrophil recruitment to the lung and subsequent IL-1β production. As the first immune cells to respond to inflammatory signals, neutrophils determine the extent of inflammatory injury(Cho, Guo et al. 2012, Kim and Luster 2015). Besides damage to the vasculature, neutrophils also induce injury in the lung epithelium, leading to alveolar damage and impaired surfactant production. Excessive recruitment and activation of neutrophils is thus negatively correlated with survival(Abraham 2003, Williams and Chambers 2014).

Recruitment of neutrophils by endothelial cells is partly controlled by expression of the adhesion molecule ICAM-1. However, we did not observe any impact of Pink1 deletion on endothelial ICAM-1 expression. We thus sought alternative explanation for how deletion of a mitochondrial protein may alter neutrophil recruitment. We found that endothelial cells release mitochondrially-encoded proteins, such as ND6, in response to inflammatory stimulus. These formylated proteins have been shown to be elevated in patient serum during sepsis, where higher amounts of circulating ND6 are correlated with higher mortality(Kwon, Suh et al. 2021). Although there are multiple possible sources of these formyl peptides, it is possible that a large proportion is derived from endothelial cells in systemic inflammatory conditions such as sepsis and LPS-induced endotoxemia. The link between mitophagy and release of mitochondrial DAMPs may on the surface appear to be an inverse one, as mitophagy generally reduces mitochondrial stress. However, several recent studies have pointed to mitochondrial stress triggering the release of mitochondrial fragments through the autophagic machinery. For instance, mitochondria in acidic lysosomes, or mitoysosomes, are released by dopaminergic neurons and astrocytes through lysosome exocytosis following Flunarizine-induced Parkinsonism(Bao, Zhou et al. 2022). Similarly, cardiomyocytes release mitochondria and mitochondrial fragments in an autophagy-dependent manner during cardiac stress(Nicolás-Ávila, Lechuga-Vieco et al. 2020).

The inflammation induced release of formylated peptides brings up an intriguing question regarding the relationship between mitochondria, which are evolutionary endosymbionts, and their eukaryotic hosts. Despite the ancient origins of our dependence on mitochondria, why are mitochondria still recognized as foreign by the immune system? Formylation of mitochondrial proteins is required for their function, as deletion of formyl-transferase leads to decreased efficiency of oxidative phosphorylation(Tucker, Hershman et al. 2011, Arguello, Köhrer et al. 2018). Perhaps inflammatory activation by formyl-peptides plays a role in host defense. The increased mortality correlated with higher ND6 levels in sepsis patients(Kwon, Suh et al. 2021) suggests that in the setting of hyperinflammation such as lung injury, the pro-inflammatory detrimental effects may override other potential host defense benefits such as amplified activation of inflammatory pathways. Mammalian immune systems may have evolved neutrophil sensing of mitochondrial formylated peptides released by the endothelium as a means of activating neutrophils even before neutrophils come into direct contact with bacteria and their formylated proteins. Such “prepping” of neutrophils when they transmigrate across the endothelial barrier by mitochondrial formylated peptides could be an essential determinant of subsequent bacterial elimination by neutrophils. However, our work suggests that there may be a need to prevent such prepping when the excessive inflammation and activation of neutrophils such as in the case of inflammatory lung injury becomes an even bigger liability than the benefits of amplified neutrophil activity.

## Materials and Methods

### Materials

Human recombinant TNFα was obtained from R&D Systems (Cat# 210-TA-020). Antibodies against Pink1 (6946S), p44/42 MAPK (Total Erk) (4695S) and phospho-p44/42 MAPK (p-Erk)(Thr202/Tyr204) (4370S) were obtained from Cell Signaling Technologies. β-Actin (SC-47778 HRP) and VE-Cadherin (sc-9989) antibodies were from SantaCruz Biotechnology. IL-1β antibody (MM425B) was obtained from Thermo Fisher Scientific. LPS, FCCP, Oligomycin and Antimycin A were obtained from Millipore Sigma. Griffonia Simplicifolia Lectin I (GSL I) Isolectin B4, DyLight® 649 was obtained from Vector Labs (Cat# DL-1208-.5). Fluorescent conjugated antibodies for flow cytometry were obtained from Biolegend - BV421-CD31 (102423), PE-CD54 (ICAM-1) (116108), APC-CD45 (103112), BV421-Ly6G (127627), PE-CD11b (101207), Bv421 Isotype Control (400259), APC Isotype Control (400612) and PE Isotype Control (400608). Purified fMLP (Formyl-Met-Leu-Phe) was obtained from Abcam (Cat# ab141806), and fMIT (Formyl-Met-Met-Tyr-Ala-Leu-Phe) was obtained from Phoenix Pharmaceuticals (Cat# 005-48).

### Cell Culture

Primary human lung microvascular endothelial cells (HLMVECs) (Cell Applications Cat# 540-05a) were cultured in flasks coated with 0.2% gelatin, using Endothelial Basal Medium 2 (Lonza Cat# CC-3156) supplemented with 10% FBS (Hyclone) and Microvascular Endothelial Growth Medium growth factor kits (Lonza Cat# CC-4147). HLMVECs between passages 5 and 9 were used for experiments. HEK293T cells were cultured in DMEM (Corning) supplemented with 10% FBS and 1% Pen-Strep (Corning). HL-60 cells were cultured in RPMI media (Corning) containing glutamine and supplemented with 10% FBS and 1% Pen-Strep and differentiated to neutrophil-like cells by supplementing media with 1.3% DMSO for 5-6 days.

### Virus Generation

Lentivirus for Mitokeima, Pink1-shRNA (Sigma Cat# TRCN0000199446) and Control-shRNA (Sigma Cat# SHC016) was generated by co-transfecting the lentiviral plasmids in HEK293T cells with VSV-G (the envelop expressing plasmid, addgene, #12259), psPax2 (the virus packaging plasmid, addgene, #12260) using JetPrime transfection reagent (Polyplus) as per the manufacturer’s protocol. Cell culture supernatant was collected 48 and 72 hours after transfection, and viral particles were precipitated using Lenti-X concentrator (Takara Bio) following the manufacturer’s protocol. HLMVECs were transduced with lentivirus in media containing 4ug/mL Polybrene (Santa Cruz Biotechnology), and expression was observed 2-4 days following infection.

### Immunofluorescence and confocal microscopy

HLMVECs expressing Mitokeima were plated on gelatin-coated glass-bottom dishes (Matek) 24 hours prior to visualization. Cells were treated as indicated and imaged live using a Zeiss Laser Scanning 710 BIG microscope equipped with a Plan-Aprochromat 63x/1.40 Oil DIC objective (Zeiss) and GaAsP and PMT detectors, at 37°C with 5% CO_2_. Mitophagy was calculated as the percentage of mitolysosome area compared to total mitochondria (mitolysosomes + cytoplasmic mitochondria). To generate movies, cells were imaged using 3x zoom for 10-20 minutes at 15 second intervals. Representative images movies were de-noised using the Noise2Void algorithm.

For ex vivo imaging, Mitokeima mice (obtained from Dr. Toren Finkel’s lab) were injected with LPS (8mg/kg, i.p.) 6 hours prior to analysis. 30 minutes prior to lung collection, anesthetized mitokeima mice aged 6-8 weeks were injected with 50 µg of Isolectin B4 (in 100 µL of PBS) retro-orbitally to stain the mouse endothelium. At the indicated time, the mouse lung was perfused with PBS and suspended in HBSS for tissue imaging. Lungs were transferred to glass-bottomed dishes in HBSS and imaged whole by confocal microscope. Images were quantified by generating a mask based on Isolectin B4 staining for the endothelium. The mask was applied to a ratiometric mask of cytoplasmic (Excitation: 488nm) to lysosomal (Excitation 560nm) mitokeima. The image was thresholded and the area of mitophagy was quantified. 10-20 fields of view were quantified per lung, 4 mouse lungs per group, from 3 independent experiments.

### Western blotting

*In vitro* samples were lysed using cell lysis buffer (50 mM HEPES pH7.5, 120 mM NaCl, 5 mM EDTA, 10 mM Na pyrophosphate, 50 mM NaF, 1 mM Na_3_VO_4,_ 1% Triton X-100) supplemented with protease (Cat# 78430, ThermoFisher Scientific) and phosphatase inhibitor (Cat# 524625, Millipore Sigma) cocktails. For mouse lung tissue samples, the post-caval lobe was flash frozen in dry ice immediately after harvesting. On thawing, tissue was homogenized (NextAdvance Bullet Blender) in lysis buffer to extract protein. Western blotting was performed as previously described, using 1:1000 dilution for all antibodies except β-actin (1:5000). Western blots were imaged using an iBright CL1500 machine (Thermo Fisher).

### Mitochondrial Stress Test

Pink1 shRNA: Control, non-targeting (Millipore Sigma, SHC016) and Pink1 (Millipore Sigma, TRCN0000199446, Seq: GAAGCCACCATGCCTACATTG) shRNA were obtained in a pLKO.1 backbone and used to generate lentivirus. 4 days following infection, HLMVECs were subjected to a Seahorse mitochondrial stress test (Agilent), following the manufacturer’s recommendations. An equal number of Pink1 and Control shRNA infected cells were plated onto a Seahorse 96 well cell culture plate. The following day, cells were washed, and media was changed to Seahorse XF Base Media, supplemented with 10mM D-Glucose (Sigma), 1mM Pyruvate (Sigma) and 2mM Glutamine (Glutamax, Gibco), containing TNFα or PBS. Mitochondrial stress test was performed 3 hours following treatment.

### fMLP and fMIT treatment of HL-60 cells

After differentiation in 1.3% DMSO, HL-60 cells were serum starved in RPMI media containing glutamine, supplemented with 0.1% FBS (RPMI-SFM) for 2 hours. 500K cells were then treated with either DMSO, 10nM fMLP or 10nM fMIT for 10 minutes. Cells were then spun down and lysed for analysis of Erk and phospho-Erk by western blot.

### Treating HL-60 cells with endothelial conditioned media

HLMVECs were plated on gelatin coated plates and allowed to grow to confluency overnight. Cells were then serum starved in EBM2 media supplemented with 0.1% FBS and treated with 10ng/µl TNFα or 10μM FCCP or left untreated for 24 hours. The following day differentiated HL-60 cells were serum starved in RPMI-SFM for 2 hours. HL-60 cells were spun down and resuspended in endothelial conditioned media that had been passed through a 0.44μm syringe filter. 10 minutes later, cells were spun down and lysed for analysis of Erk and phospho-Erk by western blot.

### Detecting ND6 in endothelial culture medium

HLMVECs were treated with either TNFα or FCCP for 24 hours. Cell culture medium was collected and filtered through 0.45µm filters to remove cell debris, and flash frozen in dry ice. ND6 was detected in samples using an ELISA kit (MyBioscource, MBS936598) as per manufacturer’s protocol, using undiluted media samples. Absorbance at 450nm was measured using a FlexStation III plate reader (Molecular Devices). Samples were normalized to untreated controls.

### Animal Procedures

All animal procedures were performed in accordance with guidelines by the UIC Animal Care and Use Committee. For retro-orbital injection, mice were anesthetized with 2% isoflurane inhalation at flow rate of 0.6 liter/min. For endpoint experiments, mice were anesthetized with intraperitoneal administration of a mixture of Ketamine (100mg/Kg), Xylazine (2mg/Kg) and acepromaxine (2 mg/Kg) in saline solution.

### DNA/liposome preparation and *in vivo* gene transfer to generate endothelial-specific Pink1 knockout mice

Cas9-VEcre mice and pGC-Pink1 sgRNAs were used to generate EC-specific Pink1 knockout mice. Cas9-VEcre were generated by breeding Cas9 (Jackson Labs 026175) and VE-Cre (Jackson Labs, 006137) mice. Mice positive for both Cas9 fl/fl and VE-Cre were used in experiments. Age-matched C57/bl6 mice (Jackson, 000664) were used as controls. Pink1 sgRNAs were designed and cloned to pGS plasmid (Genscript, #1 GCTGGTCCCGGCAAGCCGCG, #2 CAAGCGCGTGTCTGACCCAC). Liposomes were freshly made using DDAB and cholesterol as described(Orrington-Myers, Gao et al. 2006). Briefly, a mixture of DDAD and cholesterol was dissolved in chloroform, a lipid layer was formed in an evaporator (Model R-124, Rotavapor) and 5% glucose solution was added to the flask to dissolve the lipid form. Multilamellar liposomes were formed via sonication for 60 min and passed through a 0.22µm filter. 45 µg pGS-pink1 sgRNAs were gently mixed with liposome. A total volume 150 µL of the mixture was injected in either Cas9-VECre mice or C57 mice via retro-orbital injection. 4 days later, mice were treated with LPS at 8 mg/Kg body weight via intraperitoneal injection, using PBS as a vehicle control. Tissues were harvested for experiments at the indicated times. Depletion of Pink1 was confirmed by western blot.

### Evans Blue Albumin (EBA) Assay and wet/dry ratio to measure lung endothelial permeability and edema

45 minutes prior to lung collection, anesthetized mice aged 6-8 weeks were retro-orbitally injected with 100µL of 40mg/mL EBA (20 mg/Kg). At the indicated times, the lungs were perfused with 10ml of PBS at 5ml/min. The whole lungs were removed and weighed. The whole lungs in PBS were grinded and an equal volume of formamide was added to extract the EBA at 60°C overnight. The mixture was centrifuged at 5000xg for 30 min, and the absorbance of supernatants at OD620 and OD740 were measured. OD740 was used to exclude residual blood contamination, and the corrected A620 was calculated using the equation A620(corrected)=A620-1.426*A740+0.03. A calibration curve was generated using EBA, and was used to calculate the amount of EBA leaked into the lung normalized to mouse body weight.

For wet/dry ratio, the lungs were harvested without perfusion, the weight was measured. The tubes containing the lungs were dried at 60°C oven for 3 days and weighed. The ratio of wet/dry lung weight was then calculated.

### Flow Cytometry Analysis

Mouse lungs were perfused and harvested after LPS, or PBS treatment as described. Lung tissue was minced with a scissor and digested in 4mL Collagenase type 1 (1mg/mL) for 45 minutes at 37°C with gentle shaking. The tissue suspension was passed through an 18G needle 5 times, every 15 minutes during the digestion process. Following digestion, the resulting suspension was passed through a disposable 40µm strainer to remove undigested clumps. Cells were then washed with suspension buffer (PBS + 0.5% BSA + 2 mM EDTA + 4.5 mg/mL D-glucose) and resuspended in RBC Lysis Buffer (Biolegend 420301) for one minute at room temperature to remove red blood cells. Cells were washed with suspension buffer, and then blocked using TruStain FcX™ (anti-mouse CD16/32) antibody (Biolegend, 101319) in Cell Stain Buffer (Biolegend 420201) for 10 minutes at 4°C. Antibodies were added at a dilution of 1:100 in the combinations described, and cells incubated at 4°C for 30 minutes with gentle shaking. Cells were fixed in 2% Fixation buffer (Biolegend 420801) for 10 minutes, washed and then analyzed by flow cytometry. For measurement of endothelial ICAM-1 expression, ICAM-1 mean fluorescent intensity was measured in CD31+ cells. To quantify neutrophil infiltration in the lung, the percent of CD45+ Ly6G+ cells was measured by flow cytometry. Total number of cells in the lung was calculated manually using a hemocytometer to convert the percent neutrophils to the total number of neutrophils in the lung. Mean fluorescent intensity of CD11b in CD45+Ly6G+ cells was measured to indicate neutrophil activation.

### Statistical Analysis

Western blot band intensity and confocal microscopy images were quantified using ImageJ. Brightness and contrast of confocal microscopy images were adjusted for representative purposes only. Data is presented as mean ± SEM with significance levels expressed as ∗*p* < 0.05, ∗∗*p* < 0.01, ∗∗∗*p* < 0.001, and ∗∗∗∗*p* < 0.0001. All statistical analysis was performed using GraphPad Prism 8, by one-way ANOVA, with Holm-Sidak corrections for multiple comparisons, except where mentioned.

## Acknowledgements

This project was supported by NIH grant P01HL060678, 5R01HL152515-02 and 5R01HL149300-03 and AHA grant 18PRE34070092. We thank Dr. Ralf Adams for providing the VE-cadherin Cre mice. All schematic diagrams were created using Biorender.com.

## Figure Supplement Legends

**Figure 1 - figure supplement 1:**
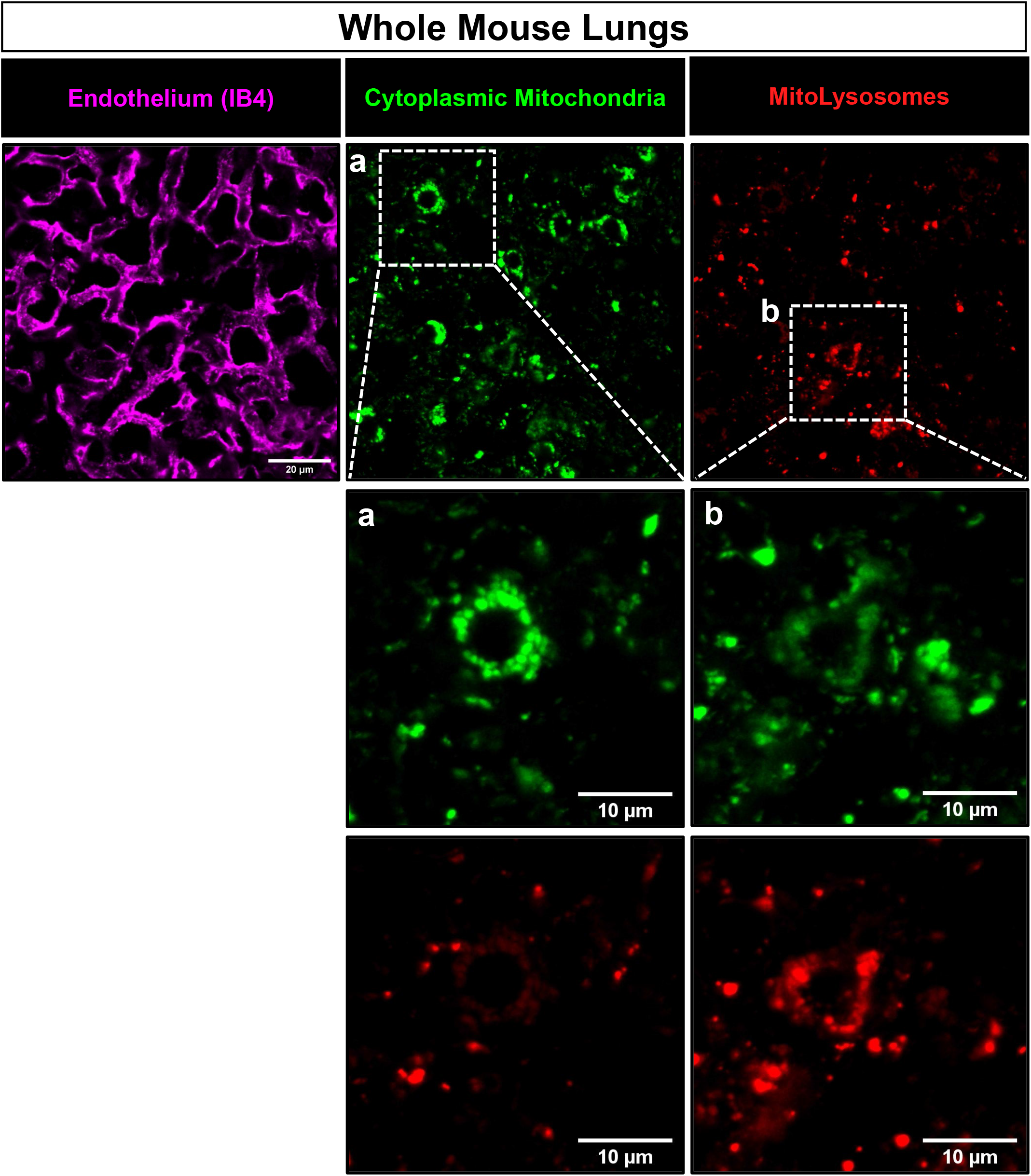
Ex vivo visualization of Mitokeima in the mouse lung. Representative images of lungs from Mitokeima mice injected with Isolectin B4 (IB4) to visualize the endothelium (magenta). Relative intensities of neutral, cytoplasmic mitochondria (green) and acidic, lysosomal mitochondria (red) are compared to identify regions of lower mitophagy (expanded inset a), and higher mitophagy (expanded inset b).

**Figure 4 - figure supplement 1:**
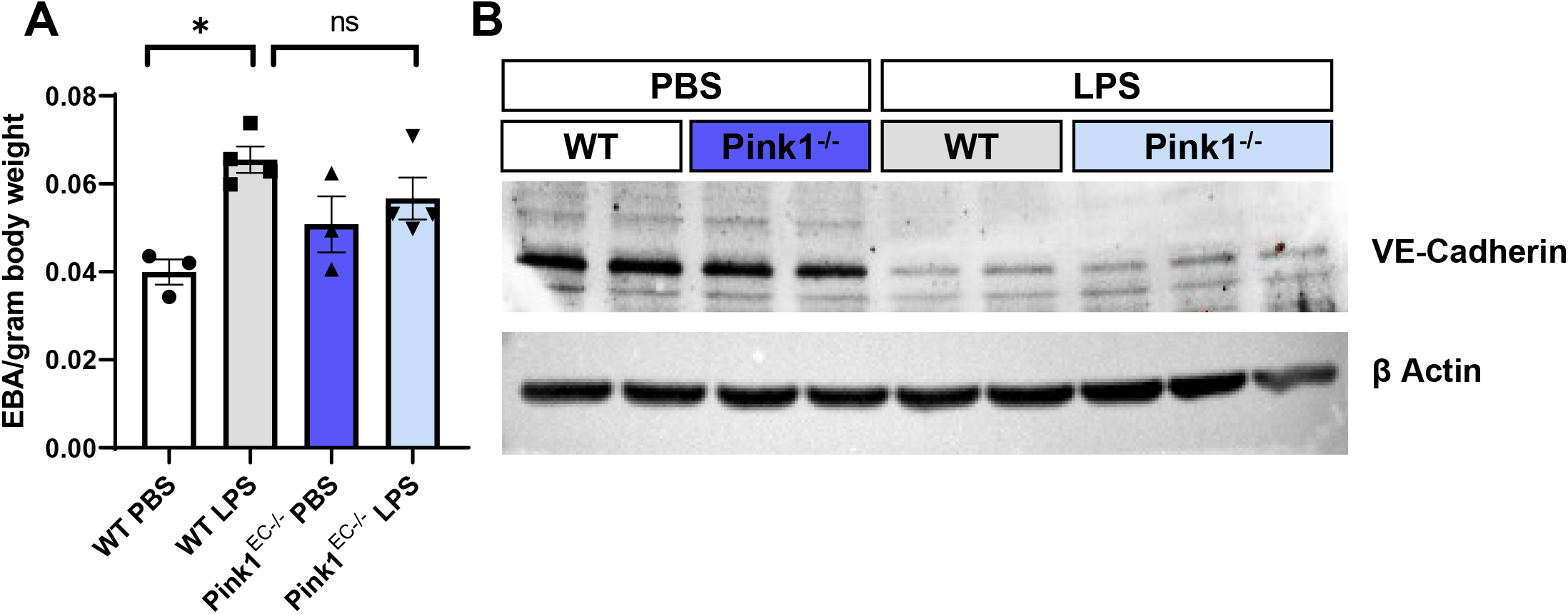
Endothelial Pink1 does not alter lung permeability. Control and Pink1^EC-/-^ mice were injected with LPS (8mg/kg), or PBS as a control. Mice were injected with Evans Blue Albumin (EBA), and lungs perfused and harvested 12-14 hours post injection. EBA in the lungs was quantified **(A)**. n=3-4 mice per group. VE-Cadherin levels were measured in lungs 6 hours following LPS injection by western blot. β-Actin was measured as a loading control. n=2-3 mice per group.

**Figure 4 - figure supplement 2:**
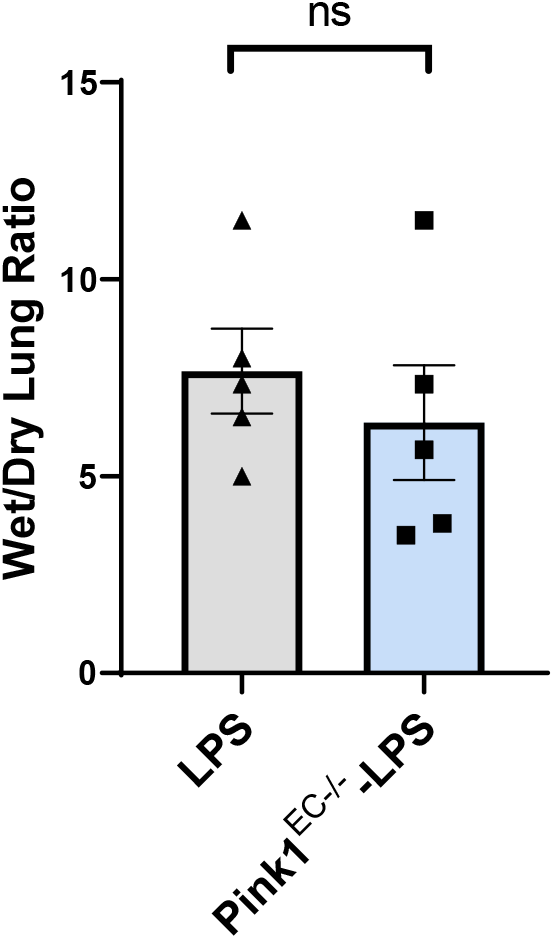
Endothelial Pink1 does not regulate lung edema. Control and Pink1^EC-/-^ mice were injected with LPS (8mg/kg), or PBS as a control. 6 hours following LPS injection, lungs were harvested and weighed. Following drying at 60°C, lungs were weighed again, and the wet-to-dry ratio was calculated as a measure of lung edema. n=5 mice per group. Statistical significance was measured by t-test.

**Figure 5 - Supplementary Figure 1.**
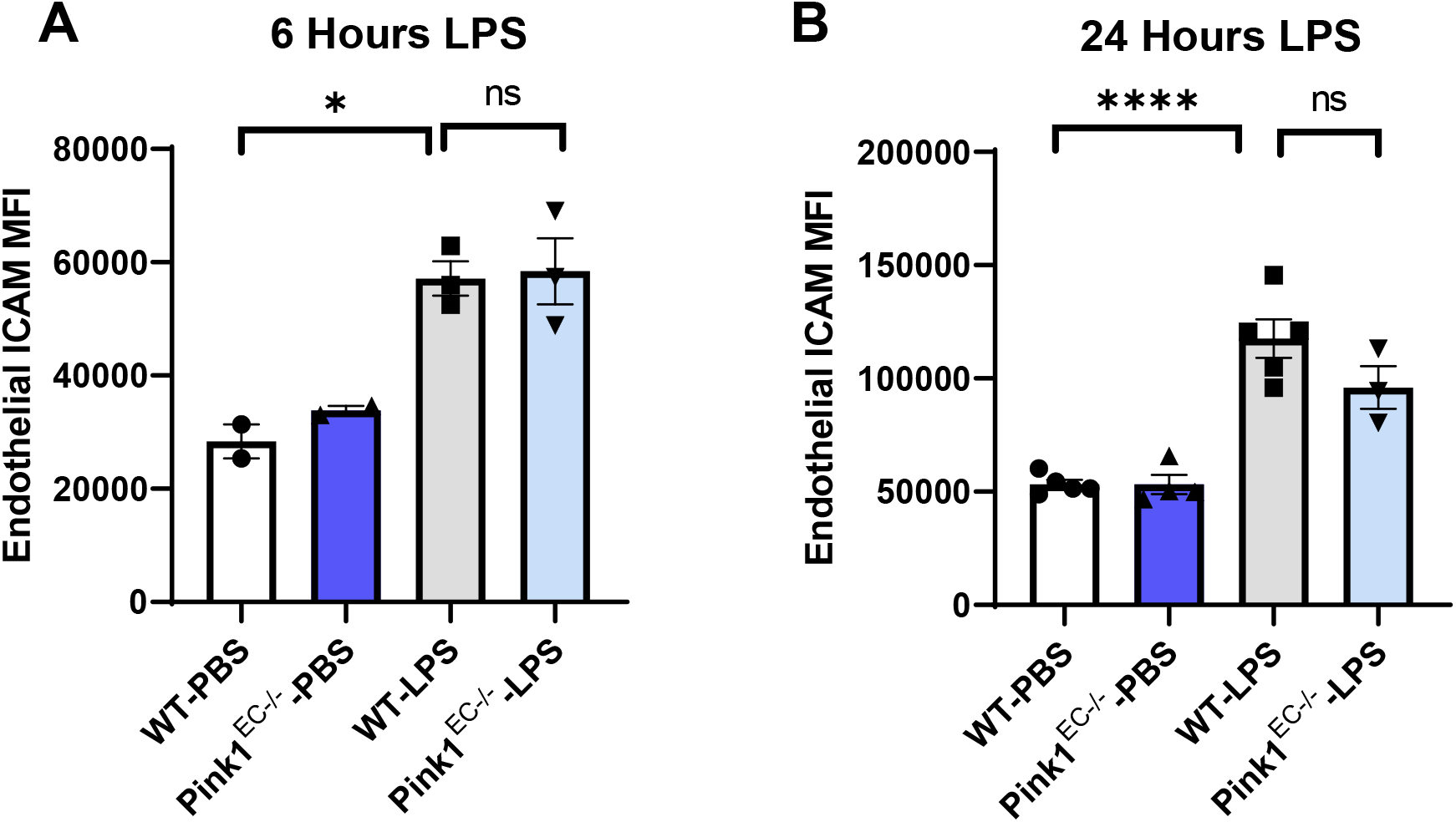

## Source Data Legends

**Figure 1 – Source Data 1**

Spreadsheet containing the original source data for the calculation of mitophagy in figure 1 D and E.

**Figure 2 – Source Data 1**

Spreadsheet containing the original source data for quantification of mitophagy in figure 2 B.

**Figure 3 – Source Data 1**

Uncropped western blot images for figures 3 A and C.

**Figure 3 - Source Data 2**

Spreadsheet containing source data for figure 3 B, D and E.

**Figure 4 – Source Data 1**

Uncropped western blot images for figure 4 B.

**Figure 4 – Source Data 2**

Spreadsheet containing source data for figure 4 C, D, supplementary figure 1 A, and supplementary figure 2.

**Figure 4 – Source Data 3**

Uncropped western blot images for figure 4 supplementary figure 1 B.

**Figure 5 – Source Data 1**

Spreadsheet containing source data for figure 5 A, B, C, D, F, and supplementary figure 1.

**Figure 5 – Source Data 2**

Uncropped western blot images for figure 5 E.

**Figure 6 – Source Data 1**

Spreadsheet containing source data for figure 6 B, D and G.

**Figure 6 – Source Data 2**

Uncropped western blot images for figure 6 C and F.

